# Genomic and functional characterization of *Pseudomonas aeruginosa*-targeting bacteriophages isolated from hospital wastewater

**DOI:** 10.1101/2021.07.08.451722

**Authors:** Hayley R. Nordstrom, Daniel R. Evans, Amanda G. Finney, Kevin J. Westbrook, Paula F. Zamora, Alina Iovleva, Mohamed H. Yassin, Jennifer M. Bomberger, Ryan K. Shields, Yohei Doi, Daria Van Tyne

## Abstract

*Pseudomonas aeruginosa* infections can be difficult to treat and new therapeutic approaches are needed. Bacteriophage therapy is a promising alternative to traditional antibiotics, but large numbers of isolated and characterized phages are lacking. We collected 23 genetically and phenotypically diverse *P. aeruginosa* isolates from people with cystic fibrosis (CF) and clinical infections, and characterized their genetic, phenotypic, and prophage diversity. We then used these isolates to screen and isolate 14 new *P. aeruginosa*-targeting phages from hospital wastewater. Phages were characterized with genome sequencing, comparative genomics, and lytic activity screening against all 23 bacterial host isolates. For four different phages, we evolved bacterial mutants that were resistant to phage infection. We then used genome sequencing and functional analysis of the resistant mutants to study their mechanisms of phage resistance as well as changes in virulence factor production and antibiotic resistance, which differed from corresponding parent bacterial isolates. Finally, we tested two phages for their ability to kill *P. aeruginosa* grown in biofilms *in vitro*, and observed that both phages reduced viable bacteria in biofilms by least one order of magnitude. One of these phages also showed activity against *P. aeruginosa* biofilms grown on CF airway epithelial cells. Overall, this study demonstrates how systematic genomic and phenotypic characterization can be deployed to develop bacteriophages as precision antibiotics.

## Introduction

The evolution of multidrug-resistant bacteria continues to outpace the development of new antimicrobials, posing a serious threat to public health. Rates of infection and mortality due to antibiotic-resistant pathogens are continuing to grow in the United States and around the world, despite efforts to curtail their spread (1, 2). Compounding the rise of multidrug-resistant bacterial infections, antibiotic development pipelines at many pharmaceutical companies have slowed or run dry (3). To help curtail this growing public health crisis, innovative approaches to antimicrobial therapy are needed.

*Pseudomonas aeruginosa* is a Gram-negative bacterium that causes a variety of infections, including bacteremia and pneumonia (4). *P. aeruginosa* chronically colonizes the airways of people with cystic fibrosis (CF), and is associated with increased morbidity and mortality in CF individuals (5). The *P. aeruginosa* species encompasses a wide breadth of genomic and phenotypic diversity, and multidrug-resistant strains often evolve during the course of prolonged antibiotic treatment (6). The success of *P. aeruginosa* as an opportunistic pathogen, its propensity for developing drug resistance, and the major threat it poses to CF patients, are compelling reasons to develop new and more effective therapies to treat *P. aeruginosa* infections.

Modern medicine is quickly approaching a “post-antibiotic” era, in which current antibiotics may no longer be effective treatments for bacterial infections due to the rampant spread of drug resistance (7). The urgent need for alternative therapies has prompted clinicians and scientists to reconsider the use of bacteriophage therapy (8), particularly for treating infections that cannot be resolved with antibiotics alone (9). Recent advances in genomics and genetic engineering have facilitated the development of phage-based therapies that have proven successful in clinical settings (9), including *P. aeruginosa* infections in CF (10). Here, we used a genetically diverse panel of 23 *P. aeruginosa* clinical isolates, collected mostly from CF patients, to isolate over a dozen distinct bacteriophages from hospital wastewater. We characterized the genomic and phenotypic diversity of the bacterial isolates and phages, including a subset of evolved phage-resistant bacterial mutants. We also tested the ability of some of the isolated phages to clear bacterial biofilms *in vitro* and *ex vivo*. These data can aid in the rational design of tailored, phage-based therapies for the treatment of *P. aeruginosa* infections.

## Materials and Methods

### Bacterial isolate collection

*P. aeruginosa* bacterial isolates were collected from patients treated at the University of Pittsburgh Medical Center (UPMC) (n=21), or were purchased from the American Type Culture Collection (ATCC) (n=2). Collection of UPMC isolates was conducted with Institutional Review Board Approval (protocol #PRO12060302). Of the UPMC isolates, 20 were collected from people with CF and one was a clinical isolate from sputum collected from a non-CF patient. Both ATCC isolates were of clinical origin. All isolates were cryopreserved in brain heart infusion (BHI) media with 16.7% glycerol and stored at −80°C.

### Hospital wastewater collection and processing

Wastewater effluent was sampled from the main sewer outflow of a Pittsburgh area hospital, at a point before the outflow joined with the municipal sewer system. A total of four samples were collected over a six-month period. Each effluent sample was centrifuged at 4,000rpm for 20 minutes to pellet solid debris, the supernatant was filtered through a 0.22-µm filter, and the sample was then concentrated by centrifugation using an Amicon 100kDa filter unit (MilliporeSigma, Burlington, MA) at 4,000rpm for 15 minutes.

### Isolation of bacteriophages and phage-resistant mutants

Lytic bacteriophages were identified with a soft agar overlay assay. Briefly, bottom agar plates were prepared containing BHI media with 1.5% agar, 1mM CaCl_2_ and 1mM MgCl_2_. Bacterial isolates were inoculated into BHI media and grown overnight at 37°C. 100µL of bacterial culture was added to a tube containing 100µL of filtered concentrated wastewater for 5-10 minutes at room temperature, and then 10mL of top agarose (BHI with 0.5% agarose, 1mM CaCl_2_, and 1mM MgCl_2_) cooled to 55°C was added and the mixture was plated onto two bottom agar plates. Plates were incubated overnight at 37°C and were examined the following day to identify lytic phage plaques. Phages were passaged by sequential picking and plating of individual plaques grown on the same bacterial isolate. Phages were picked from individual plaques using a pipette tip and were placed into 100µL of SM buffer (50mM Tris-HCl pH 7.5, 100mM NaCl, 8mM MgSO_4_) and incubated overnight at 37°C. The following day, serial 10-fold dilutions were made in SM buffer, and 3µL of each dilution was spotted onto a plate containing 5mL of top agarose mixed with 50µL of bacterial culture and layered on top of a bottom agar plate. After overnight incubation at 37°C, an individual plaque was picked and passaged again. Each phage was passaged at least twice before the generation of high-titer stocks.

To generate high-titer stocks, a single plaque was picked into 100µL of SM buffer and incubated overnight. Then, 100µL of overnight bacterial culture was added and the mixture was incubated for 5-10 minutes at room temperature, followed by addition of 10mL of top agarose and plating onto two bottom agar plates. Plates were incubated overnight at 37°C, and then plates with high plaque density were flooded with 5mL of SM buffer and incubated at 37°C for at least 1 hour to elute phages from the top agar. SM buffer was removed from each plate, pooled, spun down at 4,000rpm for 20 minutes, and filtered through a 0.22µm filter. Phage-containing lysates were extracted with 0.1 volumes of chloroform followed by 0.4 volumes of 1-octanol, and were stored at 4°C.

When phage-resistant mutants were observed in the course of preparing high-titer phage lysates, they were saved for additional characterization. Individual bacterial colonies were picked, restreaked onto BHI agar, and tested by plaque assay to confirm their resistance to the isolated phage, as well by spotting onto a lawn of the parent bacterial isolate to confirm that they were not lysogens.

### Phenotypic characterization of bacterial hosts and phage-resistant mutants

Biofilm assays were performed following a previously published protocol (11). Briefly, bacteria were first inoculated into LB media and incubated overnight at 37°C. Cultures were then diluted 1:100 into M63 media. Diluted cultures were aliquoted into vinyl 96-well plates (100µl per well) sealed, and incubated at 37°C for 24 hours. After incubation, planktonic cells were removed by inverting the plates and shaking liquid out into a sterilization tub. Plates were then submerged in water and rinsed twice to remove unattached cells. Wells were stained with 0.1% crystal violet and incubated at room temperature for 15 minutes. Plates were rinsed three times with water and shaken out vigorously, then allowed to dry completely. Crystal violet stain was solubilized with 30% acetic acid. Absorbance was read in each well at 550nm using a BioTech Synergy H1 microplate reader with GenMark software (BioTech, Winooski, VT). Two biological replicates, each containing 24 technical replicates, were run for each isolate. To test phage activity against bacteria grown in biofilms, biofilms were inoculated into 96-well plates as above and incubated for 24 hours at 37°C. Planktonic cells were removed and biofilms were washed with sterile 1×PBS using a multichannel pipettor, and either fresh M63 media or phage at 1×10^12^ PFU/mL in M63 media was added to each well. Plates were incubated for 24 hours at 37°C, then bacteria in each well were resuspended and serial 10-fold dilutions were tracked onto BHI agar plates to determine the number of colony-forming units per mL (CFU/mL) in each condition.

Extracellular protease production was measured by spotting 2.5µL of an overnight culture of each isolate grown in BHI media onto a BHI agar plate containing 10% milk. Plates were incubated overnight at 30°C and read the following day. Protease activity was detected as a clear halo surrounding the bacterial spot. Swimming motility was measured by spotting 2.5µL from an overnight bacterial culture grown in BHI media onto plates containing LB + 0.3% agar. Plates were incubated overnight at 37°C and read the following day. Swimming motility was detected as bacterial growth away from the central spot. Twitching motility was assessed by inserting a pipet tip coated in overnight bacterial culture completely through a BHI agar plate to create a small hole in the agar, and then incubating for 48 hours at 37°C. Mucoidy was measured by assessing the morphology of each isolate when grown on both LB agar and *Pseudomonas* isolation agar (PIA) plates. Each isolate was struck onto each agar type, then incubated at 37°C overnight, followed by a 48-hour incubation at room temperature.

Antibiotic susceptibility testing was performed by broth microdilution in Mueller-Hinton Broth according to the standard protocol established by the Clinical Laboratory Standards Institute (CLSI) (12). Serial two-fold dilutions of ceftazidime were tested and the minimum inhibitory concentration (MIC) was recorded as the lowest antibiotic concentration that inhibited bacterial growth by visual inspection. Pyoverdine production was measured by first growing isolates to stationary phase in LB media and then inoculating bacteria into M9 media supplemented with 20mM sodium succinate and 0.5% iron-depleted casamino acids (produced by pre-treating a 10% stock solution with 0.05g/mL Chelex-100 for one hour). Bacteria were grown overnight, then pyoverdine fluorescence was measured at excitation=400nm and emission=447nm wavelengths on a BioTech Synergy H1 microplate reader with GenMark software (BioTech, Winooski, VT). Raw fluorescence unit values were collected, background fluorescence was subtracted, and fluorescence units were normalized by OD_600_. Three biological replicates, each consisting of four technical replicates, were tested.

### Genome sequencing and analysis

Bacterial genomic DNA was extracted from 1mL overnight cultures grown in BHI media using a Qiagen DNeasy Blood and Tissue Kit (Qiagen, Germantown, MD) following the manufacturer’s protocol. Illumina sequencing libraries were prepared with a Nextera XT or Nextera kit (Illumina, San Diego, CA), and libraries were sequenced on a MiSeq using 300-bp paired-end reads, or on a NextSeq using 150-bp paired-end reads. Genomic DNA was also used to construct long-read sequencing libraries using a rapid barcoding kit (SQK-RBK004, Oxford Nanopore Technologies, Oxford, UK), and libraries were sequenced on a MinION device. Base-calling of nanopore reads was performed with Guppy. Genomes were hybrid assembled with unicycler (13), annotated with prokka (14), and were compared to one another with Roary (15). A core genome phylogenetic tree was generated using RAxML with the GTRCAT substitution model and 1000 iterations (16). Prophages were identified in each bacterial genome using PHASTER (17). Prophages of any length that were predicted to be intact or questionable by PHASTER were included. Prophage sequences were compared to one another with nucleotide BLAST, and clusters of similar prophage sequences were identified as those sharing >90% sequence coverage and >90% sequence identity.

Phage genomic DNA was extracted from 500µL of phage lysate using phenol chloroform, followed by ethanol precipitation. Briefly, 500µL phenol:chloroform:isoamyl alcohol (25:24:1) was added to each lysate, samples were vortexed and then centrifuged at 16,000 x *g* for 1 minute. The upper aqueous phase was transferred to a new tube and 500µL of chloroform was added. Samples were vortexed and centrifuged again at 16,000 x *g* for 1 minute, and the upper aqueous phase was again transferred to a new tube. Then 1µL glycogen, 0.1x volume 3M sodium acetate, and 2.5x volume 100% ethanol were added and samples were incubated overnight at −20°C. The next day samples were centrifuged at 16,000 x *g* for 30 minutes at 4°C, then the supernatant was removed and the DNA pellet was washed with 150µL 70% ethanol. DNA pellets were resuspended in 100µL nuclease-free water, and DNA was quantified with a Qubit fluorimeter (Thermo Fisher Scientific, Waltham, MA). Illumina sequencing libraries were prepared with a Nextera XT or Nextera kit (Illumina, San Diego, CA), and libraries were sequenced on a MiSeq using 300-bp paired-end reads, or on a NextSeq using 150-bp paired-end reads. Phage genomes were assembled with SPAdes v3.13.0 (18), and were annotated with prokka (14). Phage genomes were compared to one another and to other available phage genomes using BLAST (19), PHASTER (17), and Mauve (20).

### *Ex vivo* biofilm assay

Immortalized human bronchial epithelial cells isolated from a ΔF508/ΔF508 CF patient (CFBE41o- cells) (21) were cultured on Transwell inserts (Corning, Corning, NY) and grown for 7-10 days at the air-liquid interface, as described (22). Basolateral medium was replaced with minimum essential medium (MEM) supplemented with 2mM L-glutamine 24h before bacterial inoculation. *P. aeruginosa* isolate 427 was inoculated into the apical compartment as described previously (23), with the following modifications: bacteria were inoculated at a multiplicity of infection of 0.015 CFU/cell in MEM and were incubated at 37°C for 1h, followed by inoculum removal and addition of MEM supplemented with 2mM L-glutamine and 23mM L-arginine. *P. aeruginosa* phage PSA07/PB1 was added to the apical compartment 8h post-bacterial inoculation at a concentration of 8×10^6^ PFUs/mL and incubated at 37°C for 16h. Following washing of the apical and basolateral media, biofilms were collected by adding 0.1% Triton X-100 apically and centrifuged to remove soluble phages. CFUs were quantified by dilution plating onto LB agar.

### Statistical Analyses

Two-tailed *t*-tests were used to assess the significance of prophage differences between CRISPR+ and CRISPR-isolates, differences in pyoverdine production between parent and phage-resistant mutant isolates, and differences in bacterial cell density in biofilm killing experiments.

### Data Availability

Hybrid assembled bacterial host genomes were submitted to NCBI under BioProject PRJNA610040. Bacteriophage genomes were submitted to NCBI under BioProject PRJNA721956.

## Results

### *P. aeruginosa* clinical isolates used for phage screening are genetically and phenotypically diverse

To isolate bacteriophages that could be maximally useful for the treatment of *P. aeruginosa* infections, we assembled a genetically and phenotypically diverse panel of 23 *P. aeruginosa* isolates collected from clinical sources (Table S1). Two isolates were purchased from the American Type Culture Collection (ATCC), 20 isolates were collected from adults with cystic fibrosis (CF), and one isolate was collected from the sputum of a hospitalized non-CF patient. All isolates were collected from different patients. The genome of each isolate was sequenced on both the Illumina and Oxford Nanopore MinION platforms, and the resulting sequencing data were hybrid assembled (13). Over half (14/23) of the genomes were closed to a single chromosome, and the remaining nine assemblies all contained 10 or fewer contigs (Table S1). Among the 23 isolates, a total of 19 different multi-locus sequence types (STs) were identified (Table S1). A core genome phylogeny of all 23 isolates confirmed that they were highly genetically diverse (Fig. 1A). All isolates were tested for their ability to form biofilms, produce extracellular protease, exhibit swimming motility, and display a mucoid phenotype when grown on LB and *Pseudomonas* isolation agars. These phenotypes were found to be variable among the collected isolates (Fig. 1A), demonstrating that the assembled isolate panel was both genetically and phenotypically diverse.

**Figure 1.**
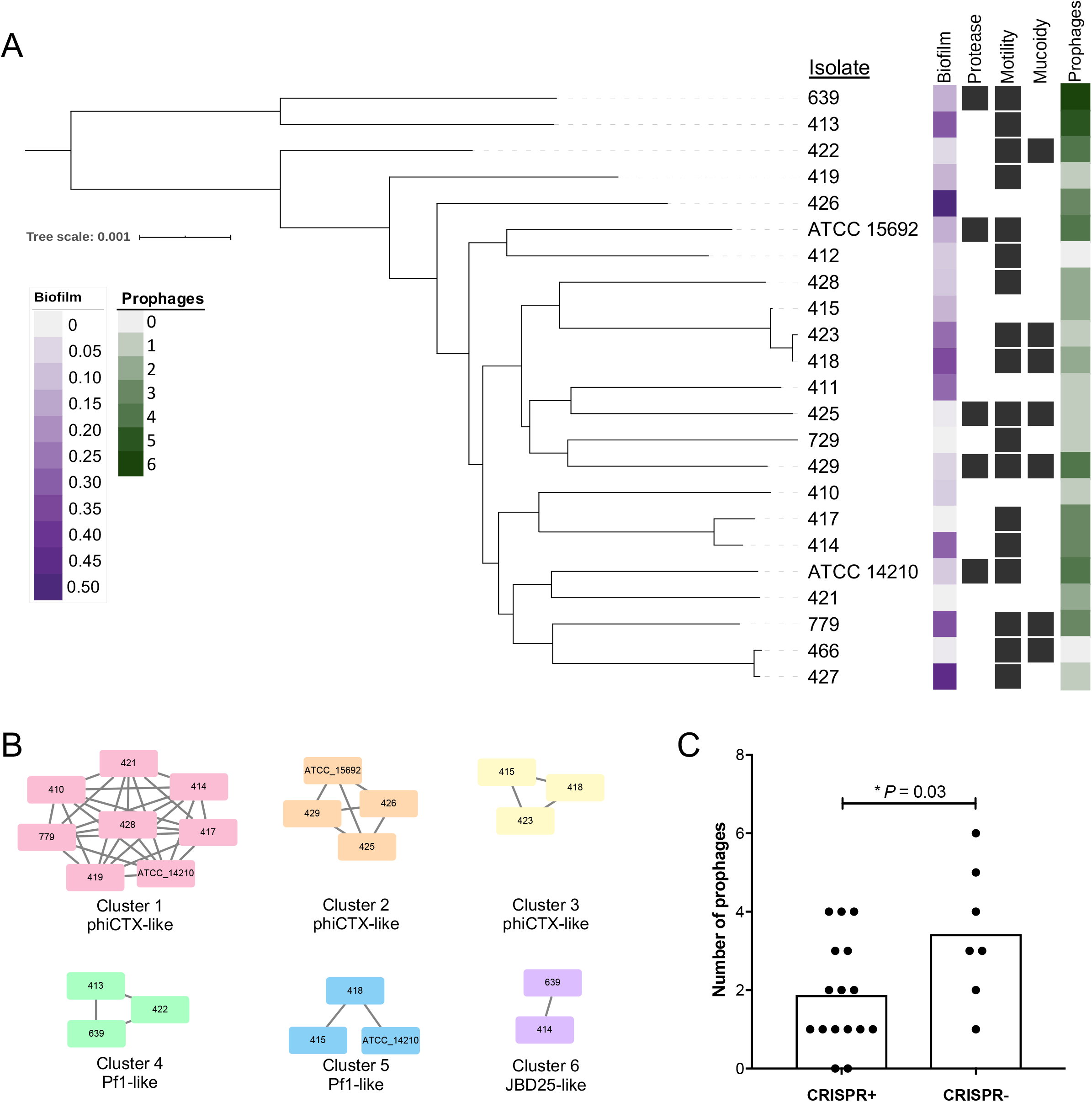
Diverse *P. aeruginosa* clinical isolates used for bacteriophage isolation and screening. (A) Core genome phylogeny of 23 *P. aeruginosa* isolates used for phage isolation. Isolates were typed for biofilm formation (measured as crystal violet staining intensity), protease production, swimming motility, mucoidy, and prophage abundance. Black squares show the presence of binary phenotypes. (B) Clusters of similar prophages found in the genomes of different *P. aeruginosa* isolates. Bacterial isolate names are listed inside the nodes of each cluster, and lines connect prophages that share >90% sequence coverage and >90% sequence identity. (C) Prophage abundance in isolates that do (CRISPR+) or do not (CRISPR-) encode functional CRISPR-Cas systems. P-value is from a two-tailed *t*-test.

We assessed the abundance and diversity of prophage sequences in the 23 *P. aeruginosa* clinical isolates we collected. The genome of each isolate was mined for prophage sequences using the PHASTER online tool (17). Between 0 and 6 prophages were found in each isolate genome (Fig. 1A, Table S2). Prophages varied in length from 5.5-74.5 Kb and in GC-content from 52.5%-66.0%. A total of 54 prophage sequences were extracted, and were compared to one another using nucleotide BLAST to assess both the nucleotide identity and coverage across all pairwise comparisons (Fig. 1B). A total of six different prophages were found to be present in more than one isolate; these included three distinct phiCTX-like phages and two Pf1-like filamentous phages. Finally, we assessed the number of prophages in isolates that were predicted to have either functional or non-functional Clustered Regularly Interspaced Short Palindromic Repeats (CRISPR) loci, based on the presence or absence of Cas enzymes in the genome of each isolate (Table S1). We found that the seven isolates predicted to have non-functional CRISPR-Cas systems had more prophages compared to isolates with intact CRISPR-Cas loci (Fig. 1C, *P*=0.03).

### *P. aeruginosa-*targeting bacteriophages isolated from hospital wastewater

We used the 23 *P. aeruginosa* isolates we collected to screen for lytic bacteriophages in wastewater effluent collected from a Pittsburgh area hospital. A total of 14 phages were isolated on 10 different *P. aeruginosa* isolates (Table 1). One additional phage, PB1, was purchased from ATCC and was propagated and characterized alongside the newly isolated phages. Because sequencing the genome of this phage revealed multiple mutations when compared to the PB1 sequence deposited in NCBI, we refer to it here as PSA07/PB1. Phages were picked and repeatedly passaged as single plaques, and were then amplified to generate high-titer stocks. Genomic DNA was extracted from each phage stock, and was sequenced on the Illumina platform. Phage genomes were found to be between 43.7-65.9 Kb in length and had GC-content ranging from 44.9%-64.5% (Table 1). Phages were compared to publicly available genomes using PHASTER and NCBI BLAST, and the predicted family and genus of each phage were determined based on similarity to previously described phages. Despite appearing to be lytic on the isolates used to propagate them, three phages (PSA04, PSA20, and PSA21) were predicted to have a lysogenic lifestyle due to the presence of annotated phage integrases. The PSA04 genome was most similar to the JBD44 lysogenic phage (24), however the homology between these two phages was not particularly high (Table 1). The PSA20 and PSA21 phage genomes showed moderate sequence similarity to the *Yuavirus* phages AN14 and LKO4, in which the putative integrase is instead believed to be a DNA primase (25). The lack of previously described lysogenic activity among *Yuavirus* phages is consistent with our observations of lytic behavior for phages PSA04, PSA20 and PSA21.

**Table 1.** Genome characteristics of *P. aeruginosa*-targeting bacteriophages. ^1^HWW = Hospital wastewater; ^2^Most similar phage based on BLAST to the NCBI nr database

| Phage | Source<sup>1</sup> | Host | Length (bp) | %GC | Predicted Family | Predicted Genus | NCBI Similar Phage (Accession)<sup>2</sup> | Coverage/Identity | BioSample |
| --- | --- | --- | --- | --- | --- | --- | --- | --- | --- |
| PSA04 | HWW | 418 | 48544 | 59.2 | *Siphoviridae* | - | JBD44 (NC_030929) | 68%/98% | SAMN18741767 |
| PSA07/PB1 | ATCC | ATCC 15692 | 65891 | 54.9 | *Myoviridae* | *Pbunavirus* | PB1 (NC_011810) | 100%/100% | SAMN18741768 |
| PSA09 | HWW | 410 | 62015 | 55.5 | *Myoviridae* | *Pbunavirus* | Pa193 (NC_050148) | 99%/93% | SAMN18741769 |
| PSA11 | HWW | ATCC 14210 | 48815 | 44.9 | *Podoviridae* | - | PA11 (NC_007808) | 97%/99% | SAMN18741770 |
| PSA13 | HWW | 427 | 45731 | 52.6 | *Podoviridae* | *Bruynoghevirus* | Pa222 (MK837011) | 99%/93% | SAMN18741771 |
| PSA16 | HWW | 466 | 45623 | 52.5 | *Podoviridae* | *Bruynoghevirus* | Pa222 (MK837011) | 98%/98% | SAMN18741772 |
| PSA20 | HWW | 639 | 62296 | 64.4 | *Siphoviridae* | *Yuavirus* | AN14 (KX198613) | 95%/98% | SAMN18741773 |
| PSA21 | HWW | 639 | 62243 | 64.5 | *Siphoviridae* | *Yuavirus* | LKO4 (NC_041934) | 96%/97% | SAMN18741774 |
| PSA25 | HWW | 426 | 64290 | 55.5 | *Myoviridae* | *Pbunavirus* | LBL3 (NC_011165) | 99%/95% | SAMN18741775 |
| PSA28 | HWW | 428 | 48440 | 58.3 | *Siphoviridae* | - | PMBT28 (MG641885) | 96%/86% | SAMN18741776 |
| PSA31 | HWW | 411 | 45505 | 52.5 | *Podoviridae* | *Bruynoghevirus* | Pa222 (MK837011) | 98%/97% | SAMN18741777 |
| PSA34 | HWW | 427 | 43749 | 52.3 | *Podoviridae* | *Bruynoghevirus* | Pa222 (MK837011) | 98%/98% | SAMN18741778 |
| PSA37 | HWW | 639 | 45506 | 52.5 | *Podoviridae* | *Bruynoghevirus* | Pa222 (MK837011) | 98%/97% | SAMN18741779 |
| PSA39 | HWW | 423 | 47030 | 64.2 | *Siphoviridae* | *Yuavirus* | LKO4 (NC_041934) | 95%/98% | SAMN18741780 |
| PSA40 | HWW | 466 | 45506 | 52.5 | *Podoviridae* | *Bruynoghevirus* | Pa222 (MK837011) | 98%/97% | SAMN18741781 |

Next, we compared the genomes of the isolated phages to one another, and to the publicly available phage genomes that were most similar to them, using nucleotide BLAST (Table 1, Fig. 2). Phages within the same genus showed varying degrees of genomic similarity with one another, and no similarity was observed across different families or genera. Six of the phages we isolated belonged to the *Bruynoghevirus* genus within the *Podoviridae* family; three of these phages (PSA31, PSA37, and PSA40) were highly similar to one another, despite having been isolated on three different *P. aeruginosa* isolates and from three different wastewater samples (Fig. S1). These data suggest that *Bruynoghevirus* phages might have been particularly abundant in the wastewater that we sampled, and that they are able to infect genetically diverse *P. aeruginosa* isolates.

**Figure 2.**
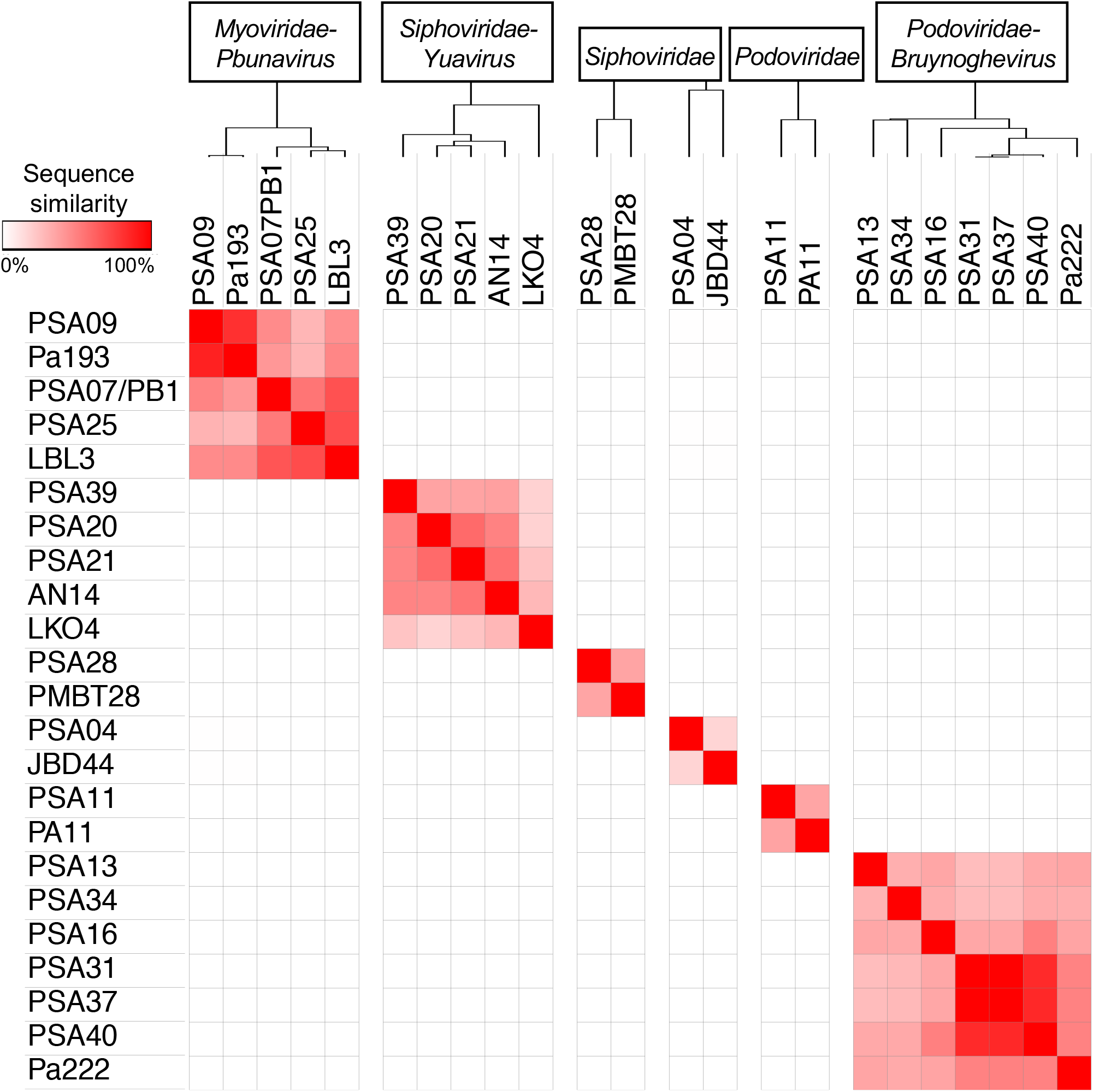
Genomic similarity among isolated *P. aeruginosa* bacteriophages and publicly available phage genomes. Phages are organized by family and genus, which are labeled at the top of the figure. Phage genomes were compared with one another using nucleotide BLAST to determine sequence coverage and nucleotide identity for each pairwise comparison. Coverage and identity values were multiplied to calculate the “sequence similarity” for each comparison. Similarity values range from 0-100%, and are shown with red shading (0% = white, 100% = red). Dendrograms at top were generated by Pearson correlation clustering of sequence similarity values across all pairwise comparisons.

### Phage susceptibility of *P. aeruginosa* isolates and bacteriophage infectivity

To examine the phage susceptibilities of our *P. aeruginosa* isolates as well as the infectivity profile of each phage, we performed a lytic activity screen of the 15 bacteriophages studied here against all 23 bacterial isolates (Fig. 3). Serial dilutions of each phage were spotted onto top agar lawns of each bacterial isolate, and individual plaques were counted to determine the titer of each phage against each isolate. Three of the *P. aeruginosa* isolates we tested (413, 414, and 729) were resistant to all phages tested, however the other 20 isolates (87% of all isolates tested) were susceptible to multiple phages belonging to different families (Fig. 3). Phage susceptibility profiles of the isolates were highly variable, with the exception of isolate pairs 418/423 and 427/466; these pairs contained isolates belonging to the same ST, which were more genetically similar to one another than to the other isolates in the study. While phages were found to infect between 9 and 19 different isolates, activity of the same phage was often variable against different isolates. For example, phage PSA07/PB1 displayed titers varying from 102 to 10^10^ PFU/mL against different *P. aeruginosa* isolates (Fig. 3). Finally, compared to *Myoviridae* and *Siphoviridae* phages, the *Podoviridae* phages we isolated were able to infect more isolates and had higher average infectivity against the isolates tested here.

**Figure 3.**
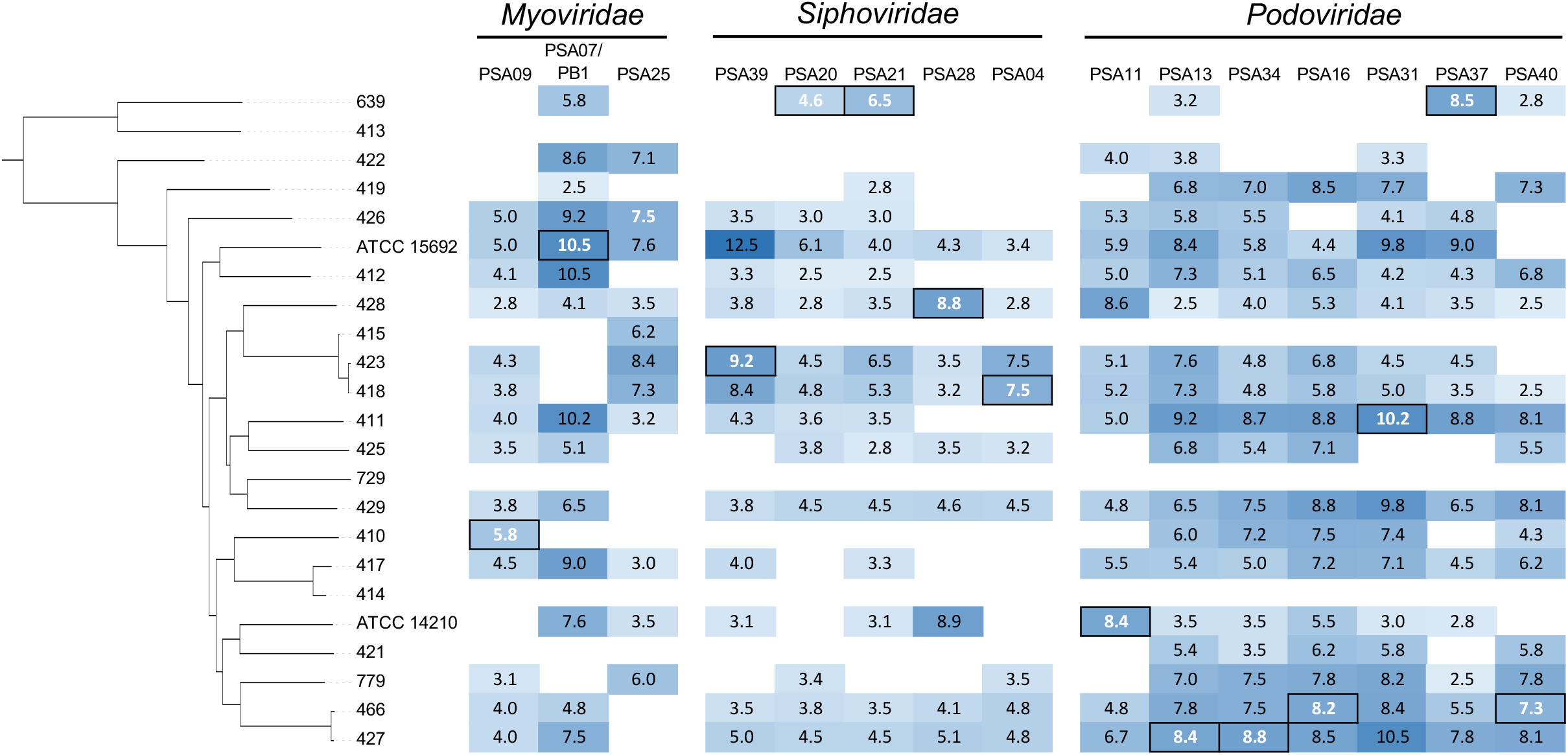
Infectivity of isolated phages against genetically diverse *P. aeruginosa* isolates. Bacterial isolates are ordered according to the core genome phylogeny in Figure 1. Infectivity is shown as the log_10_ titer (PFU/mL) of each phage against each isolate. Boxed white values indicate the *P. aeruginosa* isolate that each phage was isolated and propagated on. Blue shading corresponds to phage titer, with darker shading indicating higher titer. White shading indicates no phage activity.

### Genomic and phenotypic differences of phage-resistant mutants

During the course of phage propagation, we isolated single colonies of phage-resistant mutants for four phages: PSA09, PSA11, PSA20, and PSA34 (Fig. 4). Phage-resistant mutant isolates were tested to confirm their resistance, and were then subjected to whole-genome sequencing. Sequencing reads were mapped to the hybrid assembled genome of the corresponding phage-susceptible parent isolate, and protein-altering mutations in each resistant mutant were identified (Table 2). Each phage-resistant mutant genome encoded 2-3 protein-altering mutations. Based on the annotation of each mutated gene, we were able to identify putative phage resistance-conferring mutations in each mutant isolate genome. A phage-resistant mutant in the 639 isolate background that was resistant to phage PSA20 was found to have a Thr278Pro mutation in the Type IV pilus protein PilB (Table 2). Because Type IV pili are involved in twitching motility, we compared the twitching motility of the 639 *P. aeruginosa* parent isolate and the PSA20-resistant mutant, and found that the resistant mutant showed diminished twitching motility (Fig. 4A). Two other phage-resistant mutants raised in different *P. aeruginosa* parent isolates against different phages both encoded mutations in genes predicted to impact LPS biosynthesis, including a RfaB-like glycosyltransferase and the dTDP‑4‑dehydrorhamnose reductase RfbD (Table 2). We compared the susceptibilities of both phage-resistant mutants and their corresponding parent isolates to ceftazidime, an antibiotic that is used to treat *P. aeruginosa* infections (26). The mutants showed either four-fold or eight-fold sensitization to ceftazidime compared to their parents (Fig. 4B), indicating that the phage resistance-conferring alterations to the LPS in these mutants also increased their susceptibility to a cell wall-targeting antibiotic. A final mutant was found to carry a frameshift mutation that disrupted the coding sequence of the quorum-sensing master regulator LasR (Table 2). Because LasR is known to regulate the production of *P. aeruginosa* virulence factors, we measured the production of extracellular protease and pyoverdine in both the parent and phage-resistant mutant isolates (Fig. 4C and D). Extracellular protease production was absent and pyoverdine production was greatly diminished in the phage-resistant mutant compared to the parent isolate. Overall these data demonstrate the variability of genetic mechanisms underlying phage resistance, as well as the collateral phenotypic effects of resistance.

**Figure 4.**
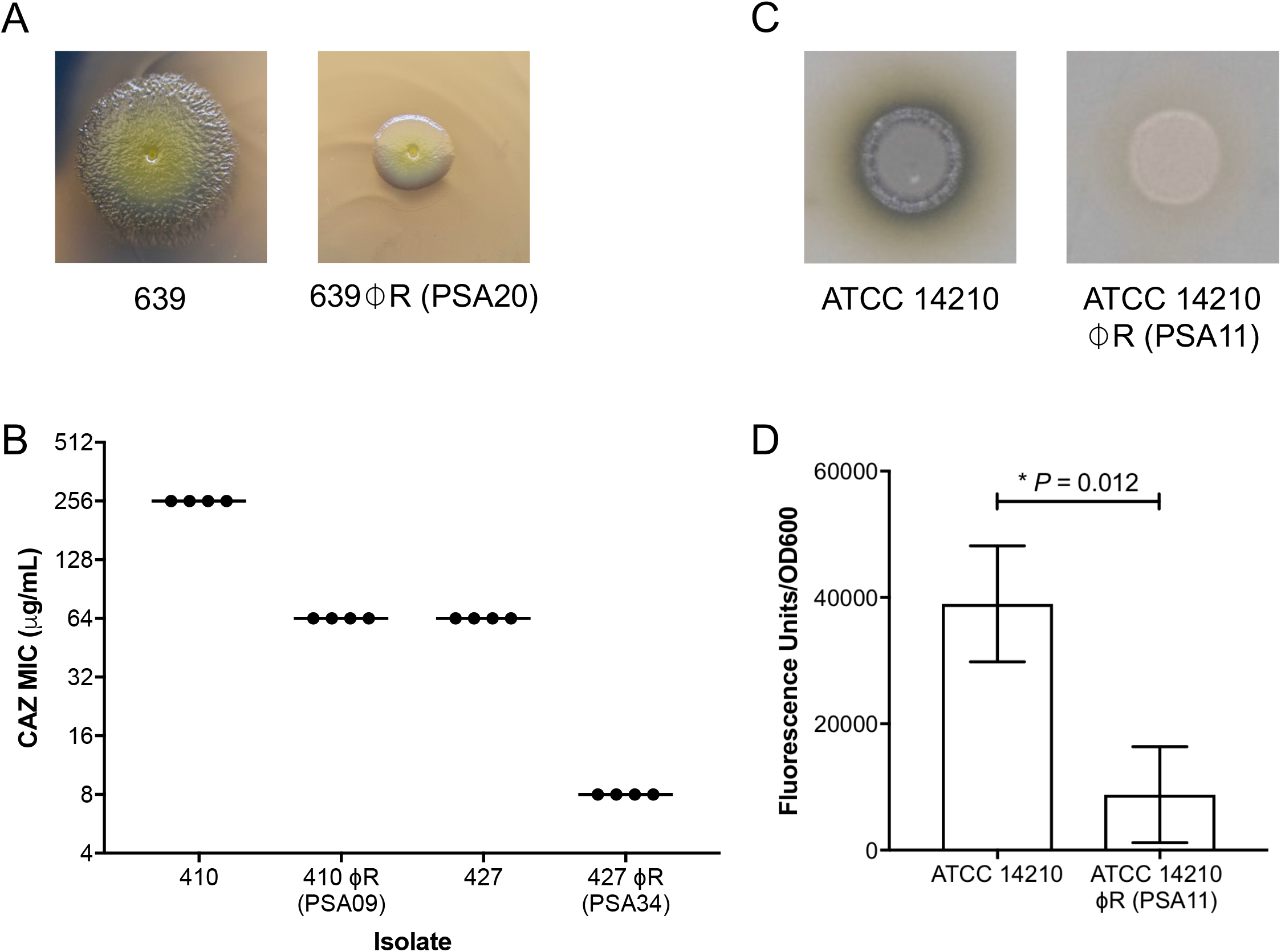
Phenotypic consequences of phage resistance. (A) Twitching motility differences between isolate 639 and 639⏀R, a phage-resistant mutant raised against phage PSA20 that harbors a mutation in the Type IV pilus protein PilB. (B) Ceftazidime (CAZ) susceptibilities of two pairs of wild type parent isolates and corresponding phage-resistant mutants, both of which harbor mutations in genes impacting LPS biosynthesis. (C) Protease production differences between isolate ATCC 14210 and ATCC 14210⏀R, a phage-resistant mutant raised against phage PSA11 that harbors a mutation in the quorum-sensing transcriptional regulator LasR. (D) Pyoverdine production quantified in ATCC 14210 and ATCC 14210⏀R. P-value is from a two-tailed *t*-test.

**Table 2.** Protein-altering mutations identified in phage-resistant *P. aeruginosa* mutants.

| Isolate | Phage | Location<sup>1</sup> | Mutation | Description |
| --- | --- | --- | --- | --- |
| 639ΦR | PSA20 | 3,133,682 | N401I | Hypothetical protein |
|  |  | 5,758,428 | T278P | Type IV pilus protein PilB |
| 410ΦR | PSA09 | 2,674,126 | V671A | 16S rRNA endonuclease CdiA |
|  |  | 2,688,288 | L5392F | 16S rRNA endonuclease CdiA |
|  |  | 5,582,688 | D279G | RfaB-like glycosyltransferase |
| 427ΦR | PSA34 | 2,909,283 | E643G | Linear gramicidin synthase subunit D IgrD |
|  |  | 5,953,386 | frameshift +TG | dTDP-4-dehydrorhamnose reductase RfbD |
| ATCC 14210ΦR | PSA11 | 3,871,727 | D304S | Cbb3-type cytochrome c oxidase subunit CcoN1 |
|  |  | 4,104,560 | frameshift +A | Transcriptional activator protein LasR |

<sup>1</sup>Genome coordinates in parent *P. aeruginosa* genome.

### Phage-mediated killing of bacterial biofilms *in vitro* and *ex vivo*

Because *P. aeruginosa* causing infections frequently grows in biofilms (27), we tested whether phages that were active against bacteria in our top agar lawn-based activity assays could also kill bacteria grown in biofilms (Fig. 5). We first tested the ability of the PSA07/PB1 and PSA34 phages to kill the 427 *P. aeruginosa* isolate grown in biofilms *in vitro*. Biofilms were grown for 24 hours, planktonic cells were removed and biofilms were washed, and then phages were applied and plates were incubated for an additional 24 hours. PSA07/PB1 treatment resulted in >100-fold bacterial killing, and PSA34 treatment resulted in >10-fold bacterial killing (Fig. 5A). Next, we tested the ability of the PSA07/PB1 phage to kill the 427 isolate grown in biofilms in association with human CF airway epithelial cells. Bacteria were incubated with epithelial cells for 8 hours, then phage was added and incubated for additional 16 hours before cell-associated bacteria were collected and quantified for viability. We found that similar to the *in vitro* assay, PSA07/PB1 treatment resulted in >100-fold bacterial killing (Fig. 5B), suggesting that phages can also kill *P. aeruginosa* grown in biofilms under conditions that more closely mimic infection in humans.

**Figure 5.**
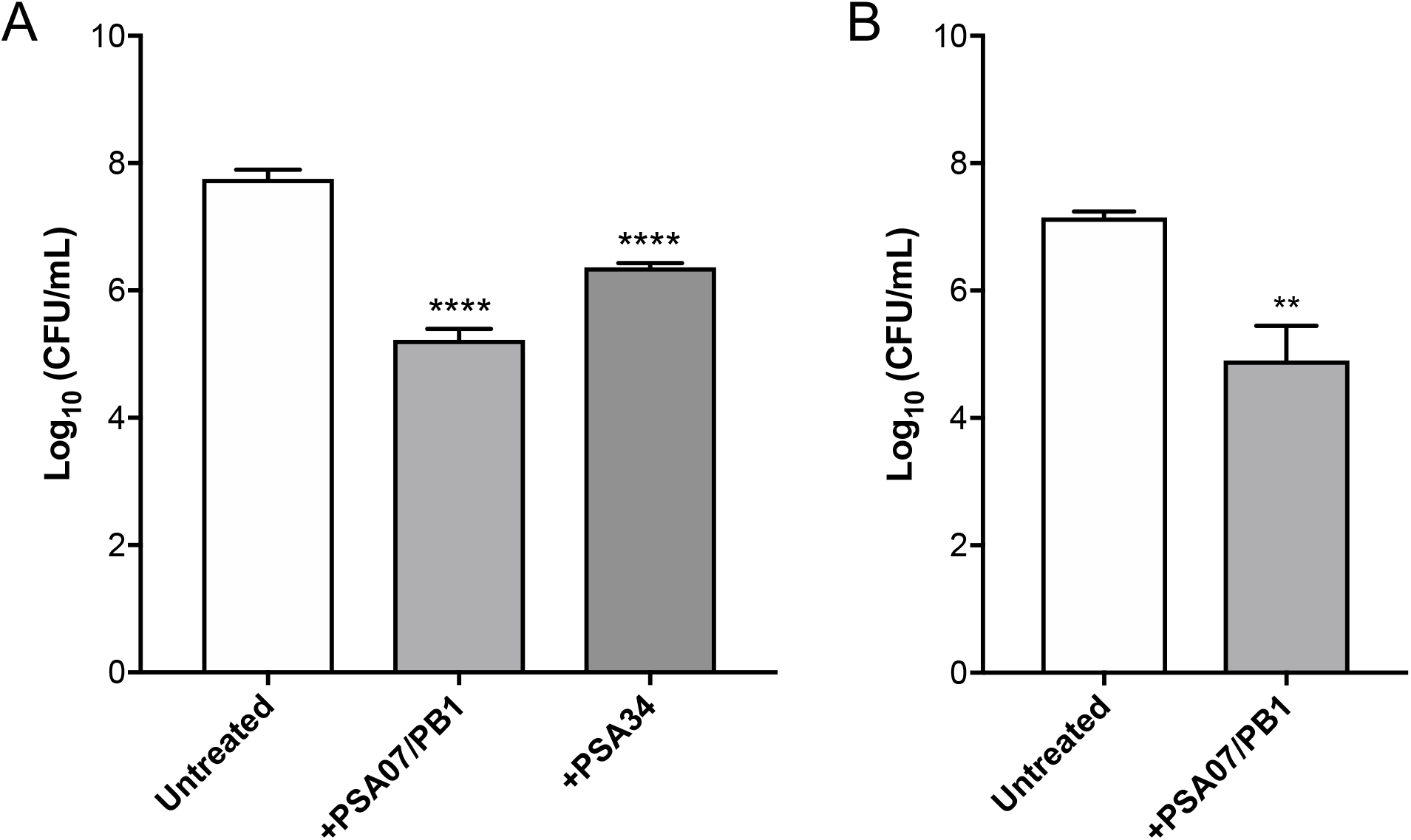
Phage-mediated killing of *P. aeruginosa* grown in biofilms *in vitro* and *ex vivo*. (A) Bacterial viability measured after *P. aeruginosa* isolate 427 biofilms were grown *in vitro* and then treated with either fresh media (Untreated), or with phages PSA07/PB1 or PSA34. (B) Viability after phage PSA07/PB1 treatment of *P. aeruginosa* isolate 427 biofilms grown on human-derived CF airway epithelial cells. Viable bacteria were quantified as CFU/mL, and phage-treated conditions were compared to the untreated condition using two-tailed *t*-tests. \*\**P* < 0.01, \*\*\*\**P* < 0.0001.

## Discussion

The objective of this study was to isolate and characterize lytic bacteriophages from hospital wastewater with activity against clinically relevant *P. aeruginosa* isolates. By screening wastewater samples against a genetically and phenotypically diverse panel of *P. aeruginosa* bacterial isolates, we were able to isolate a diverse group of *P. aeruginosa*-targeting phages representing three families: *Myoviridae*, *Siphoviridae* and *Podoviridae*. In testing our panel of *P. aeruginosa* isolates for susceptibility to the isolated phages, we found a broad range of phage activities. Additionally, our analysis of select phage-resistant mutants showed that evolving phage resistance often conferred an increase in antibiotic susceptibility or a reduction in bacterial virulence. Finally, two of the phages we isolated were able to kill *P. aeruginosa* grown in biofilms *in vitro* and *ex vivo,* suggesting that they have therapeutic utility for the treatment of *P. aeruginosa* infections.

The bacteriophages we isolated in this study were similar in terms of phage family, genus, and other genome characteristics to *P. aeruginosa*-targeting phages isolated previously (28–31). This could be due to the fact that these prior studies also isolated phages from sewage, similar to what we did in this study. Our findings, however, are in keeping with the idea that phages active against *P. aeruginosa* mirror the abundant genetic and phenotypic diversity of their hosts. While it has been noted that newly isolated phages do not often represent novel phylogenetic lineages (30), sampling and screening from more diverse sources could potentially uncover a broader range of phage genetic diversity.

While the vast majority of *P. aeruginosa* isolates we screened were susceptible to one or more of the phages we isolated, three bacterial isolates were resistant to all phages studied here. These three isolates (413, 414, and 729) were genetically distinct from one another, and no clear trends emerged to explain their resistance to phage infection. For example, they did not all have functional CRISPR-Cas systems or a higher relative abundance of prophages compared to phage-susceptible isolates. We did note that the *P. aeruginosa* 729 isolate grew very poorly, and was predicted to be a hypermutator due to a frameshift mutation in the DNA mismatch repair gene *mutS*. It is unknown whether hypermutators in *P. aeruginosa* are more resistant to phage infection; this would be a worthwhile avenue of future investigation. Nonetheless, the specific mechanism(s) conferring phage resistance in the clinical isolates we studied here remain unclear.

During the course of phage propagation, we isolated four phage-resistant *P. aeruginosa* mutants and studied them further. From whole-genome sequencing of these mutants, we identified three kinds of mutations that lead to measurable phenotypic changes. First, in the phage-resistant mutant of isolate 639, disruption of the Type IV pilus protein PilB appears to have also caused a reduction in twitching motility. Because the Type IV pilus has been previously described as a surface receptor used by *P. aeruginosa* phages for infection (32), we suspect that the phage PSA20, and also perhaps the other *Yuavirus* phages we isolated, use the Type IV pilus as a receptor for infection. Second, we identified mutations in genes affecting LPS biosynthesis in phage-resistant mutants of isolates 410 and 427. Bacterial LPS is also a well-known surface receptor used for phage infection in *P. aeruginosa* (33). Notably, we observed that the resulting phage-resistant mutants showed increased susceptibility to ceftazidime, a cell-wall targeting antibiotic. Finally, in the phage-resistant mutant of ATCC 14210, a disruption in the quorum-sensing master regulator LasR resulted in a decrease in the production of extracellular protease as well as pyoverdine, which are two prominent virulence factors in *P. aeruginosa* (34). Taken together, these findings are consistent with the notion that the development of phage resistance is often coupled with collateral effects like decreased bacterial virulence or increased antibiotic susceptibility (35). This has potentially promising implications for treatment of *P. aeruginosa* infections using phages, where a tradeoff between phage resistance and antibacterial resistance could be exploited.

When we tested whether two different *P. aeruginosa* phages could kill bacteria grown in biofilms, we observed reductions in viable bacteria upon phage treatment of biofilms both *in vitro* and *ex vivo*. While application of phage did not completely eradicate bacteria growing in the biofilms, it did substantially decrease the bacterial loads measured in both assays to similar levels to the ones obtained after antibiotic treatment (36). This finding is consistent with other studies that have also documented phage-mediated reductions in *P. aeruginosa* biofilm density *in vitro* (31, 37, 38). Here we have extended these *in vitro* findings to test the ability of phages to infect bacteria grown on human CF airway epithelial cells, a setting that more closely mimics bacterial growth in the CF airway (39, 40). Testing of phage efficacy in a context that includes eukaryotic cells is an important feature of this study. Whether and how bacteriophages interact with eukaryotic cells, and how this interaction may impact phage activity, is a focus on ongoing work by us and others (41).

This study had several limitations. Many of the phages we isolated showed variable activity, and their activity was generally diminished against isolates other than the host isolate used for their initial isolation and propagation. While we attempted to be unbiased in our phage isolation methods, we observed some redundancy in isolated phages within the *Podoviridae* family, suggesting a possible enrichment of our wastewater source with *Podoviridae* phages. Additionally, we only isolated and studied four different phage-resistant mutants, and we did not confirm that any of the resistance-associated mutations identified were indeed the cause of phage resistance, for example through genetic complementation. Finally, all work performed in this study was conducted *in vitro* or *ex vivo*, thus we are unable to conclude that any of the phages we isolated would be useful therapeutic candidates without additional testing, for example in relevant animal models of *P. aeruginosa* infection.

Taken together, the genotypic and phenotypic data presented here have promising implications for the therapeutic potential of *P. aeruginosa-*targeting bacteriophages. This study provides a valuable addition to the growing literature documenting the abundance and diversity of *P. aeruginosa* phages, and demonstrates how systematic characterization can aid in the development of phages for clinical use as precision antibiotics.

## Acknowledgements

We gratefully acknowledge all members of the Van Tyne lab, and in particular Shu-Ting Cho for helpful input during the preparation of this manuscript. We also thank Carlos Guerrero-Bustamante and Catherine Armbruster for helpful contributions to the study. Research reported in this publication was supported in part by the National Institute of Allergy and Infectious Diseases of the National Institutes of Health under Award Number UM1AI104681. The content is solely the responsibility of the authors and does not necessarily represent the official views of the National Institutes of Health. This work was also supported by grants BOMBER19R0, BOMBER21P0, and ZAMORA20F0 from the Cystic Fibrosis Foundation, and by the Department of Medicine at the University of Pittsburgh School of Medicine. The funders had no role in study design, data collection and analysis, decision to publish, or preparation of the manuscript.

## Conflicts of Interest

J.B. is a consultant for BiomX, Inc. The other authors have no relevant conflicts of interest.

---

**Figure S1.**
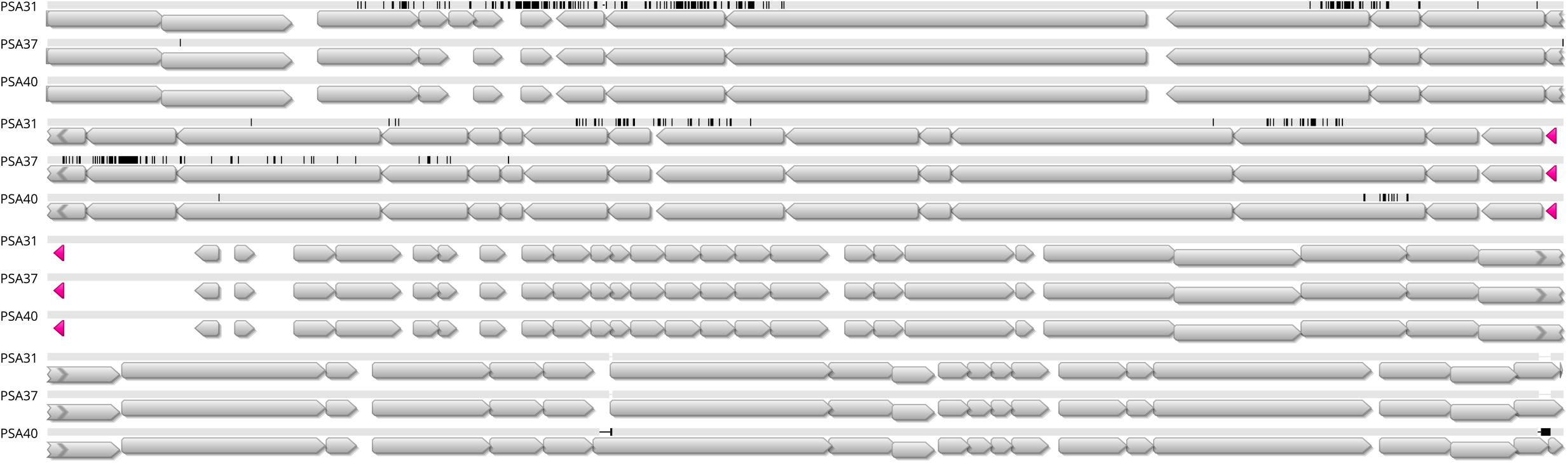
Genome sequence alignment of similar *Bruynoghevirus* phages. The genomes of phages PSA31, PSA37, and PSA40 were aligned to one another using Mauve. Grey arrows indicate coding sequences, and pink arrowheads show the location of tRNA genes. Vertical black lines show differences in nucleotide sequence between phage genomes, and horizontal black lines show differences due to nucleotide insertions or deletions.

## References

1. CDC. 2019. Antibiotic Resistance Threats in the United States, 2019 doi:http://dx.doi.org/10.15620/cdc:82532. U.S. Department of Health and Human Services, Atlanta, GA.

2. Friedman ND, Temkin E, Carmeli Y. 2016. The negative impact of antibiotic resistance. Clin Microbiol Infect 22:416–22.

3. Singer AC, Kirchhelle C, Roberts AP. 2019. Reinventing the antimicrobial pipeline in response to the global crisis of antimicrobial-resistant infections. F1000Res 8:238.

4. Moradali MF, Ghods S, Rehm BH. 2017. Pseudomonas aeruginosa Lifestyle: A Paradigm for Adaptation, Survival, and Persistence. Front Cell Infect Microbiol 7:39.

5. Høiby N, Ciofu O, Bjarnsholt T. 2010. Pseudomonas aeruginosa biofilms in cystic fibrosis. Future Microbiol 5:1663–74.

6. Winstanley C, O’Brien S, Brockhurst MA. 2016. Pseudomonas aeruginosa Evolutionary Adaptation and Diversification in Cystic Fibrosis Chronic Lung Infections. Trends Microbiol 24:327–337.

7. Alanis AJ. 2005. Resistance to antibiotics: are we in the post-antibiotic era? Arch Med Res 36:697–705.

8. Domingo-Calap P, Delgado-Martinez J. 2018. Bacteriophages: Protagonists of a Post-Antibiotic Era. Antibiotics (Basel) 7.

9. Schooley RT, Biswas B, Gill JJ, Hernandez-Morales A, Lancaster J, Lessor L, Barr JJ, Reed SL, Rohwer F, Benler S, Segall AM, Taplitz R, Smith DM, Kerr K, Kumaraswamy M, Nizet V, Lin L, McCauley MD, Strathdee SA, Benson CA, Pope RK, Leroux BM, Picel AC, Mateczun AJ, Cilwa KE, Regeimbal JM, Estrella LA, Wolfe DM, Henry MS, Quinones J, Salka S, Bishop-Lilly KA, Young R, Hamilton T. 2017. Development and Use of Personalized Bacteriophage-Based Therapeutic Cocktails To Treat a Patient with a Disseminated Resistant Acinetobacter baumannii Infection. Antimicrob Agents Chemother 61.

10. Trend S, Fonceca AM, Ditcham WG, Kicic A, Cf A. 2017. The potential of phage therapy in cystic fibrosis: Essential human-bacterial-phage interactions and delivery considerations for use in Pseudomonas aeruginosa-infected airways. J Cyst Fibros 16:663–670.

11. O’Toole GA. 2011. Microtiter dish biofilm formation assay. Journal of visualized experiments : JoVE doi:10.3791/2437:2437.

12. CLSI. 2019. Performance Standards for Antimicrobial Susceptibilty Testing, 29th ed. CLSI Supplement M100. Clinical and Laboratory Standards Institute.

13. Wick RR, Judd LM, Gorrie CL, Holt KE. 2017. Unicycler: Resolving bacterial genome assemblies from short and long sequencing reads. PLoS Comput Biol 13:e1005595.

14. Seemann T. 2014. Prokka: rapid prokaryotic genome annotation. Bioinformatics 30:2068–9.

15. Page AJ, Cummins CA, Hunt M, Wong VK, Reuter S, Holden MT, Fookes M, Falush D, Keane JA, Parkhill J. 2015. Roary: rapid large-scale prokaryote pan genome analysis. Bioinformatics 31:3691–3.

16. Stamatakis A. 2014. RAxML version 8: a tool for phylogenetic analysis and post-analysis of large phylogenies. Bioinformatics 30:1312–1313.

17. Arndt D, Grant JR, Marcu A, Sajed T, Pon A, Liang Y, Wishart DS. 2016. PHASTER: a better, faster version of the PHAST phage search tool. Nucleic Acids Res 44:W16–21.

18. Bankevich A, Nurk S, Antipov D, Gurevich AA, Dvorkin M, Kulikov AS, Lesin VM, Nikolenko SI, Pham S, Prjibelski AD, Pyshkin AV, Sirotkin AV, Vyahhi N, Tesler G, Alekseyev MA, Pevzner PA. 2012. SPAdes: a new genome assembly algorithm and its applications to single-cell sequencing. J Comput Biol 19:455–77.

19. Altschul SF, Gish W, Miller W, Myers EW, Lipman DJ. 1990. Basic local alignment search tool. J Mol Biol 215:403–10.

20. Darling AC, Mau B, Blattner FR, Perna NT. 2004. Mauve: multiple alignment of conserved genomic sequence with rearrangements. Genome Res 14:1394–403.

21. Bruscia E, Sangiuolo F, Sinibaldi P, Goncz KK, Novelli G, Gruenert DC. 2002. Isolation of CF cell lines corrected at DeltaF508-CFTR locus by SFHR-mediated targeting. Gene Ther 9:683–5.

22. Hendricks MR, Lane S, Melvin JA, Ouyang Y, Stolz DB, Williams JV, Sadovsky Y, Bomberger JM. 2021. Extracellular vesicles promote transkingdom nutrient transfer during viral-bacterial co-infection. Cell Rep 34:108672.

23. Zemke AC, Shiva S, Burns JL, Moskowitz SM, Pilewski JM, Gladwin MT, Bomberger JM. 2014. Nitrite modulates bacterial antibiotic susceptibility and biofilm formation in association with airway epithelial cells. Free Radic Biol Med 77:307–16.

24. Bondy-Denomy J, Qian J, Westra ER, Buckling A, Guttman DS, Davidson AR, Maxwell KL. 2016. Prophages mediate defense against phage infection through diverse mechanisms. The ISME Journal 10:2854–2866.

25. Evseev PVG, A. S.; Sykilinda, N. N.; Drucker, V. V.; Miroshnikov, K. A. 2020. *Pseudomonas* bacteriophage AN14 – a Baikal-borne representative of *Yuavirus*. Limnology and Freshwater Biology 5:1055–1066.

26. Nguyen L, Garcia J, Gruenberg K, MacDougall C. 2018. Multidrug-Resistant Pseudomonas Infections: Hard to Treat, But Hope on the Horizon? Curr Infect Dis Rep 20:23.

27. Maurice NM, Bedi B, Sadikot RT. 2018. Pseudomonas aeruginosa Biofilms: Host Response and Clinical Implications in Lung Infections. Am J Respir Cell Mol Biol 58:428–439.

28. Farlow J, Freyberger HR, He Y, Ward AM, Rutvisuttinunt W, Li T, Campbell R, Jacobs AC, Nikolich MP, Filippov AA. 2020. Complete Genome Sequences of 10 Phages Lytic against Multidrug-Resistant Pseudomonas aeruginosa. Microbiol Resour Announc 9.

29. Kwiatek M, Mizak L, Parasion S, Gryko R, Olender A, Niemcewicz M. 2015. Characterization of five newly isolated bacteriophages active against Pseudomonas aeruginosa clinical strains. Folia Microbiol (Praha) 60:7–14.

30. Latz S, Krüttgen A, Häfner H, Buhl EM, Ritter K, Horz HP. 2017. Differential Effect of Newly Isolated Phages Belonging to PB1-Like, phiKZ-Like and LUZ24-Like Viruses against Multi-Drug Resistant Pseudomonas aeruginosa under Varying Growth Conditions. Viruses 9.

31. Oliveira VC, Bim FL, Monteiro RM, Macedo AP, Santos ES, Silva-Lovato CH, Paranhos HFO, Melo LDR, Santos SB, Watanabe E. 2020. Identification and Characterization of New Bacteriophages to Control Multidrug-Resistant Pseudomonas aeruginosa Biofilm on Endotracheal Tubes. Front Microbiol 11:580779.

32. Bradley DE, Pitt TL. 1974. Pilus-dependence of four Pseudomonas aeruginosa bacteriophages with non-contractile tails. J Gen Virol 24:1–15.

33. Huszczynski SM, Lam JS, Khursigara CM. 2019. The Role of Pseudomonas aeruginosa Lipopolysaccharide in Bacterial Pathogenesis and Physiology. Pathogens 9.

34. Lamont IL, Beare PA, Ochsner U, Vasil AI, Vasil ML. 2002. Siderophore-mediated signaling regulates virulence factor production in Pseudomonasaeruginosa. Proc Natl Acad Sci U S A 99:7072–7.

35. Chan BK, Sistrom M, Wertz JE, Kortright KE, Narayan D, Turner PE. 2016. Phage selection restores antibiotic sensitivity in MDR Pseudomonas aeruginosa. Sci Rep 6:26717.

36. Zemke AC, Kocak BR, Bomberger JM. 2017. Sodium Nitrite Inhibits Killing of Pseudomonas aeruginosa Biofilms by Ciprofloxacin. Antimicrob Agents Chemother 61.

37. Fong SA, Drilling A, Morales S, Cornet ME, Woodworth BA, Fokkens WJ, Psaltis AJ, Vreugde S, Wormald PJ. 2017. Activity of Bacteriophages in Removing Biofilms of Pseudomonas aeruginosa Isolates from Chronic Rhinosinusitis Patients. Front Cell Infect Microbiol 7:418.

38. Fu W, Forster T, Mayer O, Curtin JJ, Lehman SM, Donlan RM. 2010. Bacteriophage cocktail for the prevention of biofilm formation by Pseudomonas aeruginosa on catheters in an in vitro model system. Antimicrob Agents Chemother 54:397–404.

39. Hendricks MR, Lashua LP, Fischer DK, Flitter BA, Eichinger KM, Durbin JE, Sarkar SN, Coyne CB, Empey KM, Bomberger JM. 2016. Respiratory syncytial virus infection enhances Pseudomonas aeruginosa biofilm growth through dysregulation of nutritional immunity. Proc Natl Acad Sci U S A 113:1642–7.

40. Cornforth DM, Diggle FL, Melvin JA, Bomberger JM, Whiteley M. 2020. Quantitative Framework for Model Evaluation in Microbiology Research Using Pseudomonas aeruginosa and Cystic Fibrosis Infection as a Test Case. mBio 11.

41. Van Belleghem JD, Dabrowska K, Vaneechoutte M, Barr JJ, Bollyky PL. 2018. Interactions between Bacteriophage, Bacteria, and the Mammalian Immune System. Viruses 11.

